# Rapidly increasing prevalence of overweight and obesity in older Ghanaian adults from 2007-2015: evidence from WHO-SAGE Waves 1 & 2

**DOI:** 10.1101/591222

**Authors:** Stella T. Lartey, Costan G. Magnussen, Lei Si, Godfred O. Boateng, Barbara de Graaff, Richard Berko Biritwum, Nadia Minicuci, Paul Kowal, Leigh Blizzard, Andrew J. Palmer

## Abstract

**Background:** Studies on changes in the prevalence and determinants of obesity in older adults living in sub-Saharan Africa are limited. We examined recent changes in obesity prevalence and associated factors for older adults in Ghana between 2007/08 and 2014/15.

**Methods:** Data on adults aged 50 years and older in Ghana were collected in the WHO SAGE Wave 1 (2007/08; n=4158) and Wave 2 (2014/15; n=1663). The weighted prevalence of obesity, overweight, normal weight and underweight, and of high central adiposity was compared in Waves 1 and 2. Multinomial and binomial logistic regressions were used to examine whether the determinants of weight status based on objectively measured body mass index and waist circumference changed between the two survey periods.

**Results:** The prevalence of obesity (Wave 1=10.2%, 95% CI: 8.9-11.7%; Wave 2=15.0%, 95% CI: 12.6-17.7%) and overweight (Wave 1=19.6%, 95% CI: 18.0-21.4%; Wave 2=24.5%, 95% CI: 21.7-27.5%) was higher in Wave 2 than Wave 1 and more than half of the population had high central adiposity (Wave 1=57.7%, 95% CI: 55.4-60.1%; Wave 2=66.9%, 95% CI: 63.7-70.0%) in both study periods. Obesity prevalence was 16% lower in males and 55% higher in females comparing Wave 1 to Wave 2. Female sex, urban residence, and high household wealth were associated with higher odds of overweight/obesity and high central adiposity. Those aged 70+ years had lower odds of obesity in both study waves. In Wave 2, females with physical activity level were more likely to be obese.

**Conclusion:** Over the 7-year period between survey waves, the population prevalence of overweight and obesity increased by 25% and 47%, respectively, while underweight reduced by 43%. These findings differed considerably by sex, which points to differential impacts of past initiatives to reduce overweight/obesity, potential high-risk groups in Ghana, and the need to increase surveillance.

## Introduction

Obesity is a significant global public health challenge because it is a major risk factor for most noncommunicable diseases (NCDs) and independently predicts overall mortality [1, 2]. Additionally, obesity is associated with poor mental health and low health-related quality of life and produces a high economic burden [3-6]. From 1980 to 2014, obesity has almost doubled among adults in most parts of the world including sub-Saharan Africa [2, 7-9]. Among urban residents in West Africa, the prevalence of obesity has doubled and has consistently increased in both men and women over a period of 15 years from 1992 to 2007 [9, 10].

In Ghana, the increasing prevalence of obesity from 1980 to 2014 among individuals aged 15-49 years has been well-documented [11-13] and the Ministry found the need to reduce obesity by 2% in the same age group within five years starting from 2008 [14]. However, little is known about the trends in the prevalence of obesity among populations of older adults aged 50 years and above. As a result of improved public health systems, faster fertility transitions, and increased life expectancy, Ghana’s population is rapidly transitioning into an ageing population [15, 16]. This is occurring concurrently with improved nutrition but increased availability of energy-dense foods and increasing sedentary lifestyle that has led to recent increases in obesity prevalence and NCDs [13, 17]. The health and productivity of those in the 50-65 years age range are important as they mentor younger colleagues and form the majority labour force for agricultural productivity that is a key sector for sustainable development and poverty reduction in Ghana [18]. Monitoring trends and identifying factors that relate to obesity provide information that allows interventions to be appropriately and effectively targeted [19, 20] but such data among older adults are lacking in Ghana. Thus, this study aimed to investigate recent changes in the prevalence of obesity among the older adult population of Ghana and identify contributing factors.

## Methods

### Study Population

Data from the World Health Organization’s (WHO) 2007/8 (Wave 1) and 2014/15 (Wave 2) Study on global AGEing and adult health (WHO-SAGE) were used. WHO-SAGE is a longitudinal study on the health and well-being of adult populations aged 50 years and older in six countries: China, Ghana, India, Mexico, Russian Federation, and South Africa [21]. Systematic replacement was used in Wave 2 to account for losses to the sample from Wave 1. In Ghana, trained SAGE teams collected individual-level data from nationally representative households of adults using a stratified, multistage cluster design. The sampling method used in both Waves 1 and 2 was based on the SAGE Wave 0 in which the primary sampling units were stratified by region and locality (urban/rural) [22-24]. Weight, height and waist circumference were measured, exempting pregnant women from weight measurements in both surveys [28].

Wave 1 had 5573 and Wave 2 had 4735 total survey respondents. The final number included in each level of analysis is shown in Figure 1. The final samples for analysis were determined after missing and biologically implausible weight, height and waist circumference measurements were excluded. Biologically implausible values were height <100cm or >250 cm, weight <30.0 kg or >250.0 kg and waist circumference < 25.0 cm or > 220 cm, and were excluded [25, 26]. Additionally, to deal with selection bias, all individuals in Wave 2 who were in Wave 1 were excluded from the analysis. Data were examined in repeated cross-sectional analysis from individuals aged 50+ years with complete responses who participated in Wave 1 (n=4158) and Wave 2 (n=1663) of the WHO-SAGE.

**Figure 1.**
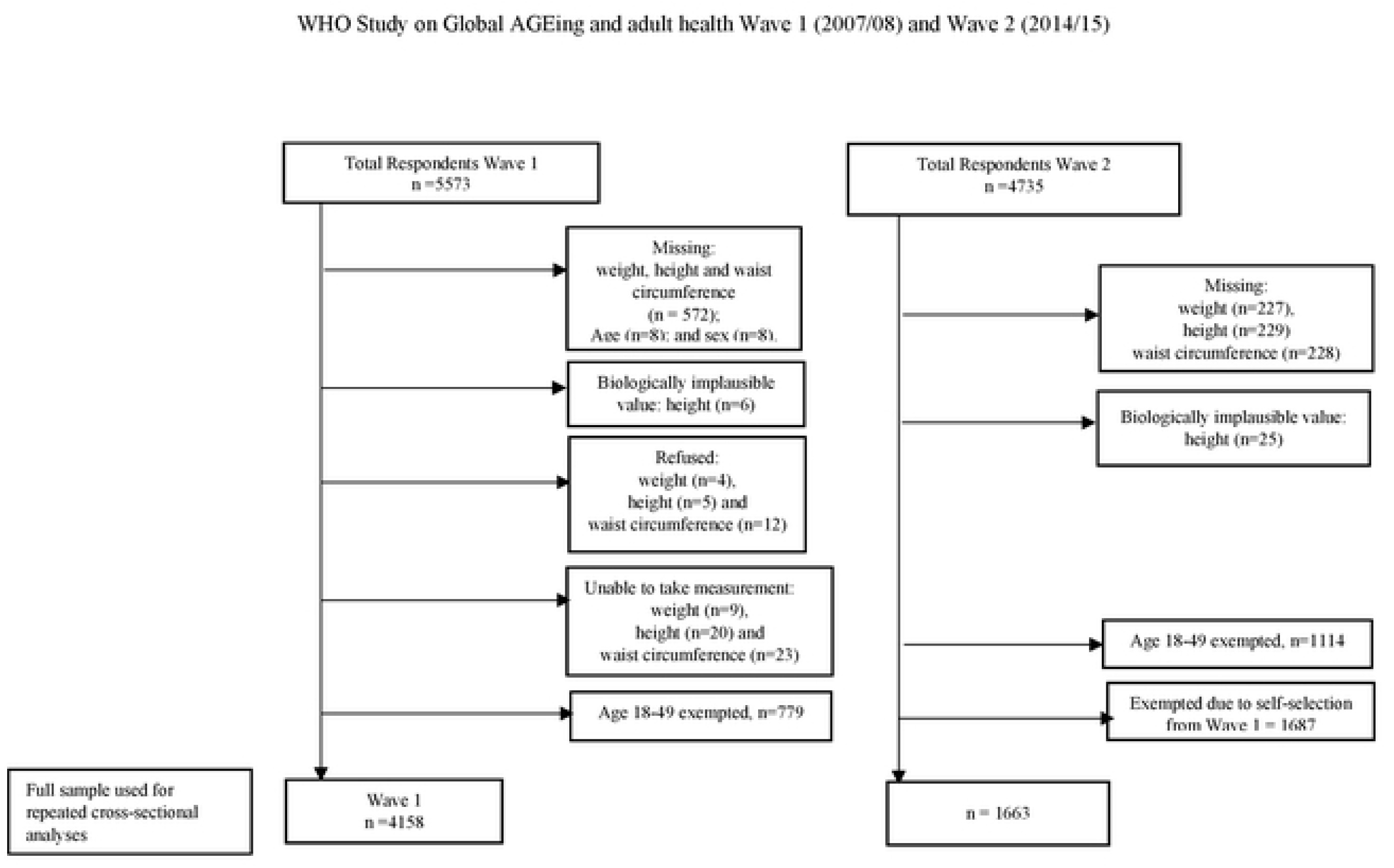
Flowchart showing samples included in the analysis. Sample extracted from Wave 1 and Wave 2 of the WHO Study on Global AGEing and adult health

## Measures

### Explanatory factors

The definition of explanatory variables used in this study followed the ecological framework developed by Scott *et al* [27] and adapts the causality continuum model for obesity in sub-Saharan Africa [28]. In their ecological theoretical framework, Scott *et al* indicated that factors influencing obesity could be situated in the distant, intermediate and proximate tiers. In the framework, three tiers overlap with distant factors found at the macro-level and influence the health outcome through the social and economic standpoint. Distant factors include globalization and urbanization, which affect factors such as lifestyle, food habits, and occupation. The intermediate factors include household and community level characteristics (e.g. household income and cultural perception about weight). Finally, the proximate factors include individual factors that directly affect the health outcome such as a genetic composition of the individual, food habits, and physical activity.

Proximate factors: These included sex, age, level of education completed and marital status. We also included smoking status, alcohol consumption, fruit and vegetable servings per day and level of physical activity per week. For estimation of age-specific prevalence, age was classified into 10-year intervals. The level of education was grouped into high education (completion of secondary/high school or higher) and low education (highest level of education completed was less than secondary or high school). Marital status was categorized as (1) single, (2) married/cohabiting and (3) divorced/separated/widowed. Respondents’ smoking status was coded as (1) “never smoked”, (2) “quitter” and (3) “current smoker”. Alcohol consumption status was coded as (1) “never”, (2) “quitter” and (3) “current drinker”. We categorized responses about fruit and vegetable intake according to international standards [29]. Respondents met the recommendation if they consumed ≥5 servings of fruits and/or vegetables per day (equivalent to 400 grams). The level of physical activity was categorised as meeting or not meeting the recommended level of total physical activity per week [30, 31].

Intermediate factors: Household wealth, a proxy for household income representing the economic status of the household [32, 33], was used as an intermediate factor. This was constructed using principal component analysis from 22 items that considered assets and the derived variable was indexed into five quintiles [34].

Distant factor: The location of participant residence (rural/ urban) was included as a distant factor in this study.

### Outcome variables

In WHO SAGE, anthropometric measurements of body weight, height, and waist circumference of respondents were taken by trained interviewers using a weighing scale, stadiometer and Gulick measuring tape following standard protocols [24]. BMI was calculated as a person’s weight in kilograms divided by the square of their height in meters and obesity was defined using cut-offs following the WHO classification. BMI was classified into four categories and weighted prevalence estimated: underweight, BMI<18.50 kg/m^2^; normal/healthy weight, BMI≥18.50<25.00 kg/m^2^; overweight, 25.00<30.00 kg/m^2^; and obese as BMI≥30.00 kg/m^2^ [35]. From the measured waist circumference, high central adiposity was determined using sub-Saharan Africa standards as waist circumference ≥81.2 cm for men, and ≥80.0 cm for women [36].

### Statistical Analysis

Accounting for the complex survey design, survey weights were used to estimate age- and sex-specific prevalence of obesity and of high central adiposity. From the cross-sectional datasets, we tested whether the frequency of distribution of the categories of BMI and central adiposity were identical in each wave using the survey Pearson’s chi-squared (*X*^*2*^) test. The absolute and percentage differences in the frequency of each category of BMI and central adiposity between the two waves were calculated for males, females, and both sexes. The categories of BMI and central adiposity were cross-tabulated against the socio-demographic and behavioral factors in Supplementary Figures 1 and 2. We fitted survey multinomial and binomial logistic models to estimate odds ratios (OR) and their 95% confidence intervals (CI) of the weight status outcome with sociodemographic and behavioural factors. The results are reported as odds ratios. Statistical analysis was performed using the complex survey model in STATA 15, and a two-tailed *p value*<0.05 was determined as statistically significant.

## Results

In total, 4158 persons (52% males) provided complete data in Wave 1 whilst 1663 new persons (41% males) did so in Wave 2. Table 1 presents the estimated frequencies of the characteristics of the study population in Waves 1 and 2. The population in Wave 2 was estimated to be around five years younger and 3.1kg heavier. Particularly for women, the proportion in the overweight/obese and high central adiposity categories were also higher in Wave 2.

**Table 1.**
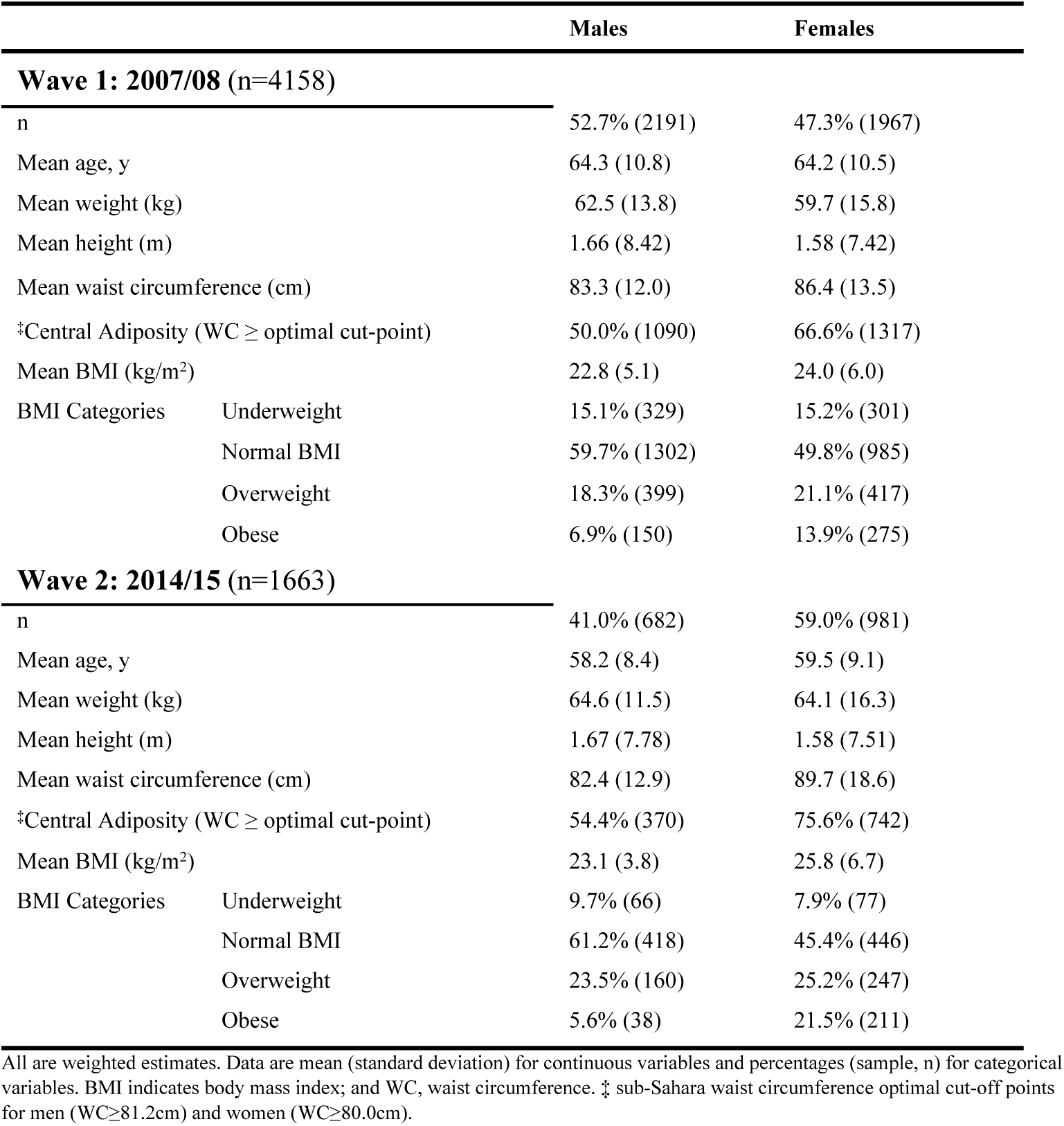
Characteristics of older people (50+ years) with complete responses in repeated cross-sectional data from WHO SAGE Waves 1 (2007/08) & 2 (2014/15)

The overall age- and sex-specific weighted prevalence of each BMI category and of high central adiposity in the repeated cross-sectional data are shown in Table 2. Relative to Wave 1, the Wave 2 prevalence of overweight (Wave 1 =19.6%; 95% CI: 18.0-21.4%; Wave 2=24.5%; 95% CI: 21.7-27.5%) and obesity (Wave 1 =10.2%; 95% CI: 8.9-11.7%; Wave 2=15.0%; 95% CI: 12.6-17.7%) was higher. Obesity increased by about 47% while overweight increased by approximately 25% in the population. More than half of the population had high central adiposity in both waves (Wave 1 =57.7%; 95% CI: 55.4-60.1%; Wave 2 =66.9%; 95% CI: 63.7-70.0%) with about a 16% increase over the 7 to 8-year period. Underweight reduced by about 43% in the population. In Wave 2, despite a decline in the prevalence of obesity for males (Wave 1=6.7%; 95% CI: 5.6-8.4%; Wave 2=5.6%; 95% CI: 3.4-8.8%), the prevalence of overweight was high for male (Wave 1=18.3%; 95% CI: 16.2-20.6%; Wave 2=23.5%; 95% CI: 18.6-29.2%); and the prevalence of both overweight (Wave 1=21.1%; 95% CI: 18.9-23.5%; Wave 2=25.2%; 95% CI: 22.4-28.2%) and obesity (Wave 1=13.9%; 95% CI: 11.8-16.4%; Wave 2=21.5%; 95% CI: 18.3-25.2%) were higher for females. The prevalence of high central adiposity was higher in Wave 2 for both males and females (Table 2).

**Table 2.** Age- and sex-specific prevalence (95% confidence interval) of body mass index (BMI) categories and high central adiposity (WC ≥ optimal cut-off point) for older people in Ghana using data from 2007/08 (n=4150) and 2014/15 (n=1663) of SAGE (All subjects with complete responses used from the repeated cross-sections).

|  |  | Underweight | Normal weight | Overweight | Obese | P Value | *High Central Adiposity | P Value |
| --- | --- | --- | --- | --- | --- | --- | --- | --- |
|  |  | *Pr (95% CI) | Pr (95% CI) | Pr (95% CI) | Pr (95% CI) |  | Pr (95% CI) |  |
| Total Sample | Wave 1 | 15.2 (13.6, 16.8) | 55.0 (52.8, 57.2) | 19.6 (18.0, 21.4) | 10.2 (8.9, 11.7) | <0.001 | 57.7 (55.4, 60.1) | <0.001 |
|  | Wave 2 | 8.6 (7.1, 10.3) | 51.9 (48.8, 55.0) | 24.5 (21.7, 27.5) | 15.0 (12.6, 17.7) |  | 66.9 (63.7, 70.0) |  |
| Absolute difference (95% CI) |  | -6.6 (-8.6, -4.4) | -3.1 (-6.7, -1.0) | 4.9 (1.6, 8.1) | 4.8 (2.0, 7.6) |  | 9.2 (5.5, 12.8) |  |
| % difference (95% CI) |  | -43.4 (-52.9, -31.2) | -5.6 (-11.8, 1.0) | 25.0 (9.1, 42.7) | 47.1 (21.6, 76.9) |  | 15.9 (10.0, 22.0) |  |
| **Males: Age, y -specific prevalence (Wave 1, n=2180; Wave 2, n=682)** |  |  |  |  |  |  |  |  |
| 50-59 | Wave 1 | 11.2 (9.0, 13.9) | 60.3 (56.3, 64.1) | 21.0 (17.9, 24.5) | 7.5 (5.8, 9.8) | 0.155 | 51.0 (46.9, 55.1) | 0.252 |
|  | Wave 2 | 6.7 (4.3, 10.5) | 61.4 (54.9, 67.5) | 25.5 (19.5, 32.7) | 6.3 (3.6, 10.8) |  | 55.0 (48.6, 61.3) |  |
| 60-69 | Wave 1 | 14.1 (11.2, 17.7) | 62.3 (57.8, 66.5) | 16.7 (13.6, 20.4) | 6.9 (4.9, 9.5) | 0.326 | 48.7 (43.8, 53.7) | 0.323 |
|  | Wave 2 | 18.2 (11.5, 27.6) | 58.0 (48.1, 67.4) | 20.3 (13.3, 29.6) | 3.5 (1.5, 8.1) |  | 54.2 (44.3, 63.8) |  |
| $\geq 70$ | Wave 1 | 21.0 (17.0, 25.5) | 56.7 (51.9, 61.4) | 16.3 (12.8, 20.4) | 6.1 (4.0, 9.1) | 0.607 | 49.2 (43.9, 54.7) | 0.158 |
|  | Wave 2 | 16.0 (9.4, 25.9) | 65.5 (54.4, 75.1) | 14.8 (8.9, 23.5) | 3.8 (1.6, 8.8) |  | 50.9 (41.0, 60.7) |  |
| Total Male sample | Wave 1 | 15.1 (13.3, 17.2) | 59.7 (57.0, 62.3) | 18.3 (16.2, 20.6) | 6.7 (5.6, 8.4) | 0.022 | 49.8 (46.8, 52.8) | 0.117 |
|  | Wave 2 | 9.7 (7.4, 12.8) | 61.2 (56.2, 66.1) | 23.5 (18.6, 29.2) | 5.6 (3.4, 8.8) |  | 54.4 (49.2, 59.5) |  |
| Absolute difference (95% CI) |  | -5.4 (-8.6, 2.2) | 1.5 (-4.0, 7.1) | 5.2 (-0.5, 10.8) | -1.1 (-4.4, 1.7) |  | 4.6 (-1.1, 10.4) |  |
| % difference (95% CI) |  | -35.8 (-52.2, -13.4) | 2.5 (-7.0, 13.3) | 28.4 (2.4, 64.1) | -16.4 (-51.0, 33.0) |  | 9.2 (-2.3, 22.1) |  |
| **Females: Age, y -specific prevalence (Wave 1, n=1398; Wave 2, n=981)** |  |  |  |  |  |  |  |  |
| 50-59 | Wave 1 | 9.7 (7.1, 13.2) | 45.7 (40.8, 50.6) | 24.7 (20.7, 29.1) | 20.0 (16.0, 24.6) | 0.012 | 70.6 (66.5, 74.5) | 0.014 |
|  | Wave 2 | 5.2 (3.7, 7.2) | 42.6 (38.2, 47.2) | 26.3 (22.7, 30.3) | 25.8 (21.6, 30.6) |  | 77.7 (73.3, 81.5) |  |
| 60-69 | Wave 1 | 12.2 (9.2, 16.1) | 53.3 (48.1, 58.5) | 22.1 (18.5, 26.2) | 12.3 (9.8, 15.5) | 0.125 | 69.6 (64.1, 74.6) | 0.389 |
|  | Wave 2 | 8.5 (5.3, 13.2) | 51.1 (43.6, 58.5) | 21.5 (16.6, 27.3) | 18.9 (13.0, 26.8) |  | 73.5 (65.7, 80.1) |  |
| $\geq 70$ | Wave 1 | 24.3 (13.0, 19.3) | 51.8 (47.4, 56.1) | 15.9 (13.0, 19.3) | 8.0 (5.8, 10.8) | 0.042 | 59.1 (54.1, 63.9) | 0.017 |
|  | Wave 2 | 17.9 (12.6, 24.7) | 49.1 (41.4, 56.8) | 25.5 (18.5, 33.9) | 7.6 (4.3, 13.1) |  | 69.8 (62.1, 76.6) |  |
| Total Female Sample | Wave 1 | 15.2 (13.1, 17.5) | 49.8 (46.7, 52.8) | 21.1 (18.9, 23.5) | 13.9 (11.8, 16.4) | <0.001 | 66.6 (63.6, 69.4) | <0.001 |
|  | Wave 2 | 7.9 (7.9, 10.8) | 45.4 (41.7, 49.1) | 25.2 (22.4, 28.2) | 21.5 (18.3, 25.2) |  | 75.6 (71.9, 78.9) |  |
| Absolute difference (95% CI) |  | -7.3 (-10.0, -4.7) | -4.4 (-8.9, 0.1) | 4.1 (0.3, 7.9) | 7.6 (3.7, 11.5) |  | 9.0 (4.7, 13.3) |  |
| % difference (95% CI) |  | -48.0 (-59.4, -34.0) | -8.8 (-16.7, -0.1) | 19.4 (2.5, 39.1) | 54.7 (26.8, 88.3) |  | 13.5 (7.7, 19.6) |  |

\*Pr (95% CI) means prevalence (95% confidence intervals). All are weighted estimates. \*High central adiposity measured using sub-Saharan high waist circumference cut-off point for men (WC ≥ 81.2cm) and women (WC ≥ 80.0cm)

The distribution of prevalence of the BMI categories by socio-demographic and behavioral factors in Wave 1 and Wave 2 are shown in Supplementary Figures 1 and 2. In general, the prevalence of obesity was high in individuals who resided in urban areas and those from households with high/higher wealth status in both Waves. While obesity was high in Wave 1 among those with high education, in Wave 2 it was rather high among both males and females with low education. Obesity was low among females who met the recommended physical activity level in both Waves.

The Univariable analyses results are presented in Table 3. In the multivariable analyses in Wave 1, being female (overweight: OR=1.47, 95% CI: 1.15-1.89; obesity: OR=2.49, 95% CI: 1.74-3.55; and high central adiposity: OR=2.03, 95% CI: 1.64-2.51), living in an urban area (obesity: OR=2.01, 95% CI: 1.41-2.86; and high central adiposity: OR=1.51, 95% CI: 1.19-1.92) and those from households with moderate, higher or higher wealth (overweight: OR=4.42, 95% CI: 2.84-6.89; obesity: OR=4.91, 95% CI: 2.41-10.17; and high central adiposity: OR=3.70, 95% CI: 1.22-6.85) were associated with higher odds of overweight, obesity and central adiposity (Table 4). However, being in age group 70+ years (overweight: OR=0.70, 95% CI: 0.54-0.90; obesity: OR=0.54, 95% CI: 0.36-0.80), currently smoking status (overweight: OR=0.54, 95% CI: 0.35-0.83; and high central adiposity: OR=0.69, 95% CI: 0.53-0.90), and current alcohol drinking (overweight: OR=0.64, 95% CI: 0.50-0.82) were associated with lower odds of overweight/obesity. Also, being in the 70+ years age group (underweight: OR=2.21, 95% CI: 1.73-2.84) and currently smoking status (underweight: OR=1.63, 95% CI: 1.18-2.25) were associated with higher odds of underweight while the odds of underweight was low among those with high/higher household wealth. In Wave 2, most associations found in Wave 1 remained the same with minimal changes in magnitude.

**Table 3.** Univariable regressions: Factor associated with BMI categories and central adiposity in year WHO-SAGE Wave 1 (2007/08) and wave 2 (2014/15)

|  |  | Year 2007/08 |  |  | Year 2014/15 |  |  | Year 2007/08<br>High Central<br>Adiposity | Year 2014/15<br>High Central<br>Adiposity |
| --- | --- | --- | --- | --- | --- | --- | --- | --- | --- |
|  |  | Underweight | Overweight | Obese | Underweight | Overweight | Obese |  |  |
|  |  | OR (95% CI) | OR (95% CI) | OR (95% CI) | OR (95% CI) | OR (95% CI) | OR (95% CI) | OR (95% CI) | OR (95% CI) |
| Age Groups, y | 50-59 | 1.00 | 1.00 | 1.00 | 1.00 | 1.00 | 1.00 | 1.00 | 1.00 |
|  | 60-69 | 1.17 (0.88, 1.53) | 0.78 (0.62, 0.98) * | 0.66 (0.48, 0.89) ** | 1.95 (1.21, 3.16) ** | 0.77 (0.55, 1.07) | 0.73 (0.45, 1.18) | 0.94 (0.79, 1.12) | 0.94 (0.70, 1.26) |
|  | ≥ 70 | 2.11 (1.66, 2.69) *** | 0.70, (0.56, 0.87) ** | 0.51 (0.36, 0.74) *** | 2.76 (1.70, 4.49) *** | 0.79 (0.53, 1.18) | 0.35 (0.20, 0.60) *** | 0.77 (0.65, 0.93) ** | 0.84 (0.63, 1.12) |
| Sex | Male | 1.00 | 1.00 | 1.00 | 1.00 | 1.00 | 1.00 | 1.00 | 1.00 |
|  | Female | 1.20 (0.97, 1.49) | 1.39 (1.13, 1.68) ** | 2.42 (1.84, 3.20) *** | 1.09 (0.75, 1.58) | 1.45 (1.05, 2.00) * | 4.22 (3.14, 8.70) *** | 2.01 (1.72, 2.34) *** | 2.59 (1.97, 3.40) *** |
| Educational<br>Status | Low | 1.00 | 1.00 | 1.00 | 1.00 | 1.00 | 1.00 | 1.00 | 1.00 |
|  | High | 0.80 (0.62, 1.03) | 1.38 (1.10, 1.74) | 2.14 (1.61, 2.84) | 2.43 (1.42, 4.14) ** | 0.98 (0.67, 1.43) | 0.80 (0.53, 1.21) | 1.57 (1.30, 1.91) | 0.76 (0.55, 1.06) |
| Marital status | Single | 1.00 | 1.00 | 1.00 | 1.00 | 1.00 | 1.00 | 1.00 | 1.00 |
|  | Married/ cohabiting | 0.56 (0.27, 1.17) | 1.65 (0.78, 3.49) | 1.11 (0.41, 2.98) | 1.14 (0.21, 3.31) | 1.84 (0.69, 4.85) | 0.87 (0.28, 2.73) | 1.07 (0.64, 1.80) | 0.98 (0.49, 1.96) |
|  | Widow/divorcee | 0.79 (0.39, 1.60) | 1.59 (0.75, 3.38) | 1.72 (0.69, 4.31) | 1.65 (0.29, 4.45) | 1.18 (0.44, 3.18) | 0.95 (0.29, 3.08) | 1.48 (0.89, 2.46) | 0.98 (0.50, 1.93) |
| Location | Rural | 1.00 | 1.00 | 1.00 | 1.00 | 1.00 | 1.00 | 1.00 | 1.00 |
|  | Urban | 0.62 (0.47, 0.82) ** | 2.32 (1.82, 2.97) *** | 4.06 (2.56, 7.20) *** | 0.47 (0.29, 0.78) ** | 1.89 (1.36, 2.61) *** | 3.47 (2.14, 5.62) *** | 2.61 (2.05, 3.32) *** | 2.35 (1.73, 3.19) *** |
| Wealth Index | Lowest | 1.00 | 1.00 | 1.00 | 1.00 | 1.00 | 1.00 | 1.00 | 1.00 |
|  | Low | 0.84 (0.60, 1.16) | 1.73 (1.20, 2.51) ** | 2.76 (1.29, 5.89) ** | 1.08 (0.65, 1.79) | 1.13 (0.64, 1.99) | 1.43 (0.43, 3.73) | 1.60 (1.27, 2.02) *** | 1.49 (0.99, 2.23) |
|  | Moderate | 0.70 (0.52, 0.95) * | 2.56 (1.76, 3.72) *** | 3.25 (2.04, 6.82) *** | 1.43 (0.78, 2.63) | 2.00 (1.18, 3.40) * | 3.72 (1.98, 6.87) ** | 2.06 (1.59, 2.66) *** | 2.24 (1.44, 3.48) *** |
|  | High | 0.70 (0.48, 1.01) | 2.66 (1.81, 3.91) *** | 3.92 (2.32, 7.44) *** | 0.88 (0.46, 1.65) | 2.56 (1.53, 4.28) *** | 4.26 (2.81, 8.25) *** | 2.40 (1.84, 3.12) *** | 2.78 (1.75, 4.41) *** |
|  | Higher | 0.34 (0.21, 0.54) *** | 4.26 (2.33, 7.05) *** | 4.94 (2.46, 9.54) *** | 0.28 (0.31, 0.60) ** | 3.27 (1.92, 5.58) *** | 5.17 (3.07, 10.10) *** | 3.95 (2.80, 6.90) *** | 3.84 (2.37, 6.22) *** |
| Smoking status | Never Smoked | 1.00 | 1.00 | 1.00 | 1.00 | 1.00 | 1.00 | 1.00 | 1.00 |
|  | Quitter | 0.99 (0.71, 1.40) | 0.77 (0.57, 1.05) | 0.37 (0.23, 0.60) *** | 1.58 (0.50, 3.01) | 1.08 (0.40, 2.92) | 1.84 (0.31, 4.98) | 0.97 (0.76, 1.25) | 1.02 (0.36, 2.91) |
|  | Currently smoke | 1.78 (1.33, 2.38) *** | 0.32 (0.22, 0.47) *** | 0.34 (0.17, 0.68) ** | 1.87 (0.88, 3.97) | 0.18 (0.07, 0.48) ** | 0.18 (0.04, 0.81) * | 0.67 (0.55, 0.80) *** | 0.23 (0.13, 0.41) *** |
| Alcohol<br>Consumption<br>Status | Never drunk<br>alcohol | 1.00 | 1.00 | 1.00 | 1.00 | 1.00 | 1.00 | 1.00 | 1.00 |
|  | Quitter | 0.88 (0.64, 1.21) | 0.76 (0.56, 1.03) | 1.00 (0.70, 1.44) | 0.90 (0.43, 1.91) | 0.54 (0.30, 0.98) * | 1.57 (0.82, 3.01) | 0.59 (0.47, 0.73) *** | 1.84 (1.03, 3.27) * |
|  | Currently Drinks | 1.18 (0.93, 1.51) | 0.57 (0.45, 0.71) *** | 0.75 (0.55, 1.02) | 0.94 (0.58, 1.51) | 0.64 (0.42, 0.97) * | 0.69 (0.38, 1.26) | 0.37 (0.28, 0.47) *** | 0.69 (0.49, 0.96) * |
| Fruit &<br>Vegetable<br>Intake | Below requirement | 1.00 | 1.00 | 1.00 | 1.00 | 1.00 | 1.00 | 1.00 | 1.00 |
|  | Met requirement | 0.86 (0.66, 1.12) | 1.17 (0.95, 1.42) | 1.32 (1.02, 1.72) * | 0.57 (0.39, 0.84) ** | 1.23 (0.91, 1.67) | 1.32 (0.91, 1.92) | 1.61 (1.32, 1.96) *** | 1.27 (0.95, 1.71) |
| Total Physical<br>Activity | Below requirement | 1.00 | 1.00 | 1.00 | 1.00 | 1.00 | 1.00 | 1.00 | 1.00 |
|  | Met requirement | 0.85 (0.67, 1.08) | 0.88 (0.71, 1.10) | 0.64 (0.47, 0.87) ** | 1.05 (0.72, 1.54) | 0.85 (0.62, 1.17) | 0.81 (0.55, 1.19) | 0.76 (0.63, 0.92) ** | 0.84 (0.63, 1.11) |

\* $p < 0.05$ , \*\* $p < 0.01$ , \*\*\* $p < 0.001$ ; Fruit & Vegetable Intake (serving per day); Total Physical Activity (Minutes per week); Year is the year for completion of data collection

**Table 4.** Multivariable regressions: predictors of overweight, obesity and central adiposity in year WHO-SAGE Wave 1 (2007/08) and wave 2 (2014/15)

|  |  | Year 2007/08 |  |  | Year 2014/15 |  |  | Year 2007/08<br>High Central<br>Adiposity | Year 2014/15<br>High Central<br>Adiposity |
| --- | --- | --- | --- | --- | --- | --- | --- | --- | --- |
|  |  | Underweight | Overweight | Obese | Underweight | Overweight | Obese |  |  |
|  |  | OR (95% CI) | OR (95% CI) | OR (95% CI) | OR (95% CI) | OR (95% CI) | OR (95% CI) | OR (95% CI) | OR (95% CI) |
| Age Groups, y | 50-59 | 1.00 | 1.00 | 1.00 | 1.00 | 1.00 | 1.00 | 1.00 | 1.00 |
|  | 60-69 | 1.17 (0.88, 1.55) | 0.79 (0.61, 1.10) | 0.67 (0.49, 0.91) * | 1.94 (1.19, 3.18) ** | 0.77 (0.55, 1.09) | 0.59 (0.36, 0.97) * | 1.01 (0.83, 1.23) | 0.93 (0.71, 1.22) |
|  | ≥ 70 | 2.21 (1.73, 2.84) *** | 0.70, (0.54, 0.90) ** | 0.54 (0.36, 0.80) ** | 2.45 (1.49, 4.04) *** | 0.87 (0.57, 1.31) | 0.34 (0.18, 0.62) ** | 0.86 (0.70, 1.07) | 0.94 (0.68, 1.29) |
| Sex | Male | 1.00 | 1.00 | 1.00 | 1.00 | 1.00 | 1.00 | 1.00 | 1.00 |
|  | Female | 1.23 (0.88, 1.71) | 1.47 (1.15, 1.89) ** | 2.49 (1.74, 3.55) *** | 0.81 (0.52, 1.26) | 1.70 (1.16, 2.49) ** | 4.65 (2.65, 9.09) *** | 2.03 (1.64, 2.51) *** | 3.36 (2.42, 4.68) *** |
| Educational<br>Status | Low | 1.00 | 1.00 | 1.00 | 1.00 | 1.00 | 1.00 | 1.00 | 1.00 |
|  | High | 1.34 (1.02, 1.78) * | 0.97 (0.73, 1.28) | 1.15 (0.83, 1.60) | 1.86 (1.00, 3.42) | 1.22 (0.80, 1.86) | 0.94 (0.61, 1.46) | 1.17 (0.93, 1.47) | 0.74 (0.49, 1.14) |
| Marital status | Single | 1.00 | 1.00 | 1.00 | 1.00 | 1.00 | 1.00 | 1.00 | 1.00 |
|  | Married/ cohabiting | 0.60 (0.29, 1.23) | 2.00 (0.92, 4.35) | 1.60 (0.64, 3.96) | 1.17 (0.17, 2.93) | 2.25 (0.81, 4.21) | 1.61 (0.63, 4.11) | 1.25 (0.69, 2.25) | 1.54 (0.69, 3.43) |
|  | Widow/divorcee | 0.74 (0.37, 1.08) | 1.50 (0.68, 3.31) | 1.65 (0.69, 3.95) | 1.39 (0.19, 3.94) | 1.20 (0.44, 3.28) | 1.11 (0.43, 2.84) | 1.16 (0.67, 2.02) | 1.10 (0.46, 2.23) |
| Location | Rural | 1.00 | 1.00 | 1.00 | 1.00 | 1.00 | 1.00 | 1.00 | 1.00 |
|  | Urban | 0.79 (0.57, 1.08) | 1.44 (1.08, 1.92) * | 2.01 (1.41, 2.86) *** | 0.53 (0.32, 0.87) * | 1.38 (0.96, 1.98) | 1.73 (1.03, 2.89) * | 1.51 (1.19, 1.92) ** | 1.91 (1.33, 2.74) ** |
| Wealth Index | Lowest | 1.00 | 1.00 | 1.00 | 1.00 | 1.00 | 1.00 | 1.00 | 1.00 |
|  | Low | 0.87 (0.62, 1.21) | 1.52 (1.05, 2.21) * | 2.40 (1.13, 5.13) * | 1.12 (0.65, 1.91) | 1.13 (0.66, 1.95) | 1.53 (0.45, 3.21) | 1.31 (1.02, 1.69) * | 1.43 (0.93, 2.20) |
|  | Moderate | 0.77 (0.51, 1.17) | 2.01 (1.33, 3.04) ** | 3.17 (1.50, 6.68) ** | 1.91 (1.04, 3.50) * | 2.03 (1.16, 3.55) * | 3.17 (1.87, 6.31) ** | 1.56 (1.18, 2.07) ** | 1.66 (1.02, 2.69) * |
|  | High | 0.71 (0.51, 0.97) * | 2.15 (1.46, 3.16) *** | 3.79 (1.96, 7.93) *** | 1.20 (0.68, 2.67) | 2.57 (1.50, 4.42) ** | 3.76 (2.27, 7.28) *** | 1.57 (1.16, 2.11) ** | 2.33 (1.42, 3.81) ** |
|  | Higher | 0.40 (0.23, 0.68) ** | 4.42 (2.84, 6.89) *** | 4.91 (2.41, 10.17) *** | 0.52 (0.22, 1.21) | 3.29 (1.89, 5.74) *** | 4.67 (2.39, 8.53) *** | 3.70 (3.22, 6.85) *** | 2.45 (1.39, 4.32) ** |
| Smoking status | Never Smoked | 1.00 | 1.00 | 1.00 | 1.00 | 1.00 | 1.00 | 1.00 | 1.00 |
|  | Quitter | 1.08 (0.74, 1.57) | 0.92 (0.68, 1.26) | 0.53 (0.30, 0.92) * | 1.68 (0.50, 3.64) | 1.73 (0.61, 4.95) | 3.48 (0.86, 6.19) | 0.77 (0.59, 0.10) | 1.06 (0.43, 2.60) |
|  | Currently smoke | 1.63 (1.18, 2.25) ** | 0.54 (0.35, 0.83) ** | 0.88 (0.46, 1.69) | 1.66 (0.69, 4.00) | 0.31 (0.11, 0.90) * | 0.48 (0.10, 2.23) | 0.69 (0.53, 0.90) ** | 0.44 (0.22, 0.88) * |
| Alcohol<br>Consumption<br>Status | Never drunk<br>alcohol | 1.00 | 1.00 | 1.00 | 1.00 | 1.00 | 1.00 | 1.00 | 1.00 |
|  | Quitter | 0.88 (0.63, 1.23) | 0.71 (0.51, 0.97) * | 0.85 (0.57, 1.29) | 0.68 (0.32, 1.43) | 0.56 (0.30, 1.06) | 1.47 (0.64, 3.38) | 0.96 (0.74, 1.24) | 2.52 (1.38, 4.59) ** |
|  | Currently Drinks | 1.19 (0.93, 1.54) | 0.64 (0.50, 0.82) *** | 0.95 (0.68, 1.31) | 0.91 (0.56, 1.49) | 0.78 (0.48, 1.27) | 1.21 (0.70, 2.08) | 0.86 (0.72, 1.03) | 1.11 (0.78, 1.58) |
| Fruit &<br>Vegetable<br>Intake | Below requirement | 1.00 | 1.00 | 1.00 | 1.00 | 1.00 | 1.00 | 1.00 | 1.00 |
|  | Met requirement | 0.98 (0.75, 1.27) | 1.05 (0.85, 1.29) | 1.16 (0.90, 1.49) | 0.57 (0.37, 0.86) ** | 1.16 (0.85, 1.60) | 1.25 (0.82, 1.90) | 1.51 (1.23, 1.84) *** | 1.27 (0.92, 1.72) |
| Total Physical<br>Activity | Below requirement | 1.00 | 1.00 | 1.00 | 1.00 | 1.00 | 1.00 | 1.00 | 1.00 |
|  | Met requirement | 0.81 (0.64, 1.02) | 1.01 (0.80, 1.27) | 0.78 (0.57, 1.08) | 1.12 (0.73, 1.71) | 0.82 (0.59, 1.14) | 0.72 (0.49, 1.08) | 0.82 (0.67, 0.99) * | 0.89 (0.66, 1.20) |

\* $p < 0.05$ , \*\* $p < 0.01$ , \*\*\* $p < 0.001$ ; Fruit & Vegetable Intake (serving per day); Total Physical Activity (Minutes per week); Year is the year for completion of data collection

An interaction term between the age categories and sex in a multivariable regression showed that the product of age and sex were not significantly associated with the BMI categories for all age groups in both Waves 1 and 2 except for females between the age of 50-59 years in whom there was a significant association for high odds of obesity (OR=2.67; 95% CI: 1.34-5.30) in Wave 1. The same interaction using central adiposity as the outcome also revealed significantly higher odds of high central adiposity among females between ages 50-59 years (OR=1.69; 95% CI: 1.10-2.58) and 60-69 years (OR=1.54; 95% CI: 1.06-2.25) only in Wave 1. We also examined whether respondents sex modified the association between physical activities and the BMI categories as well as central adiposity. No significant associations were found in Wave 1. However, in Wave 2, not meeting the recommended physical activity among females was associated with higher odds of obesity (OR=3.23; 95% CI: 1.13-6.23) and central adiposity (OR=2.19; 95% CI: 1.32-3.63).

## Discussion

This study estimated changes in the prevalence and determinants of BMI and central adiposity in the older adult population of Ghana between year and 2007/08 and 2014/15. We found that over the 7 to 8-year period the prevalence of obesity had increased by 47%, overweight by 25%, but underweight reduced by 43%. However, we found heterogeneity by sex with females showing a 55% increase in the prevalence of obesity compared with a 28% reduction among males over the same period. Being female, living in an urban area and having high household wealth were associated with higher odds of obesity/high central adiposity while those aged 70+ years was associated with lower odds of obesity. Additionally, in Wave 2, not meeting the recommended physical activity among females was associated with higher odds of obesity and central adiposity.

Our findings show that while the prevalence of underweight reduced over the period, overweight, obesity and central adiposity increased over the same period. The increased prevalence of overweight, obesity and central adiposity could have a negative public health implication as obesity buttresses the increasing burden of NCDs in most LMICs. The increase in the prevalence of overweight and obesity in our population over the 7-8year period was mainly driven by females. Even though most previous studies in sub-Saharan Africa were cross-sectional studies, most of the studies found a higher prevalence of overweight/obesity in the female population compared to their male counterparts [9, 13, 37, 38]. This phenomenon has been attributed to the body preference of most females [9, 38-40]. However, a current study of African-Americans suggested that being male with West-African ancestry genes could be protective against obesity specifically, high central adiposity and hence could be a reason for the male/female disparities [41]. Additionally, high odds of overweight/obesity among African females in many parts of sub-Saharan Africa and elsewhere, have also been largely attributed to general cultural preference in which overweight/obesity is regarded as a source of beauty and a sign of affluence [37, 40, 42]. However, in Ghana, this could also be attributed to generally low levels of physical activities in the population which could be due to the loss or lack of open and safe places for physical activities [37, 43]. Thus, this ties in with our finding that low physical activities in females was associated with higher odds of obesity and central adiposity suggesting that promotion of physical activity may support efforts to reduce obesity in older females in Ghana as in other population [20, 44, 45].

While obesity declined in Wave 2 in all age groups in males, overweight and obesity prevalence in both waves was high in females in all age groups with a decline noted after age 59 years. The low obesity prevalence observed in males may not necessarily mean that males aged 50+ years are protected. Rather it could be attributed to lifestyle changes males within this age group might have adopted, and male’s perception of an ideal body weight resulting in a reduction in their predisposition to weight gain. A study found that in sub-Saharan Africa perception about ideal body image and childhood circumstances of the different sexes as well as female’s dominance in the control of household food spending might significantly explain the sex differentials in obesity prevalence [46]. The study found that women perceived an ideal body image of a women to be larger while men perceived men’s image to be otherwise. Additionally, while women who were nutritionally deprived during childhood were more likely to be obese during adulthood, this not the same for males [46]. While obesity prevalence was found to generally increase with age in some developed countries, in most developing countries, it was found to peak around age 50 years, then decline afterward [47]. In this study a decline in obesity prevalence rates was found after age 59 years in females. Such trends may demand deliberate attention as those aged over 50 years supply the majority of the labour force for agricultural productivity, which has played a major role in sustainable development and poverty reduction in Ghana [18]. Increased obesity prevalence in this age group may lead to higher NCDs and consequently increase overall mortality as well as increasing the medical cost of care [6]. This can further lead to increased absenteeism from work and reduced work productivity, negatively impacting the economy [6, 48].

Urban residency was also associated with higher odds of overweight/obesity and central adiposity in both Waves. Residing in an urban area has shown a similar association with obesity in previous studies in Africa and elsewhere [9, 49, 50]. Urban residency is mostly associated with changes in diets and food availability, increased dependence on mechanized transportation, especially in older people coupled with an increasingly sedentary lifestyle and the loss or lack of open and safe places for physical activities [27, 51]. Dietary transition from nutritious foods to the consumption of easily accessible cheap calorie-dense foods as well as longer hours spent on buses and in cars in traffic may have been key drivers of increasing obesity prevalence in urban areas [43, 51, 52], and this is likely to be the same in Ghana. Older people, especially those over the ages 65 years, tend to rely heavily on their children and grandchildren for activities of daily living [53]. These activities include food preparation and timely food supply.

These children, who may be busy on the labour market, may be restricted in their ability to prepare home-made foods for them. They may, therefore, resort to purchasing calorie-dense fast-foods. It is also possible that the reliance on others for food sources/preparation amongst those aged over 70 years may contribute to loss of weight, which agrees with our finding that those aged 70+ years had higher odds of underweight.

The finding that high household wealth is associated with higher odds of overweight/obesity and central adiposity in both Waves in this population concurs with other findings in most LMICs [37, 54]. Household wealth, a proxy for household income, was expected to have a positive impact resulting in good health outcomes for the individuals within a household. However, in the case of Ghana, individuals from high-income households tend to be obese. Our findings support Philipson and Posner’s [54] argument that income has a positive impact on weight in less-developed economies; however, as economic development improves, this relationship tends to be negative in the long term. Increased household wealth could potentially contribute to changes in food preference, increased food consumption and poor choices regarding dietary intake.

As part of lifestyle factors, our finding that smoking was associated with lower odds of overweight has been found in previous studies where current smokers were less likely to have increased body weight compared to those who have never smoked before [55]. In Wave 1 quitting smoking was associated with lower odds of obesity however in Wave 2, currently smoking was rather associated with lower odds of overweight which agrees with previous findings in a randomized control trial [56]. As our findings are from repeated cross-sectional studies, we are unable to confer causality with further studies necessary to confirm the strength and direction of the association.

This study has several strengths. First, missing from the extant literature is a study that tracks trends in obesity prevalence among older adults in Ghana. We used the most current data to measure prevalence, changes in prevalence and factors associated with overweight/obesity among older adults about whom little is known in sub-Saharan Africa. Second, the prevalence estimates will be useful in establishing and predicting the future economic burden of obesity on health in this population. Third, the inclusion of waist circumference, a marker of central adiposity and a prime marker of cardiometabolic diseases, provides added confirmation of our findings. Finally, the use of a population-based data that is representative of the older adult population, and uses objectively measured rather than self-reported weight, height, and waist circumference to determine obesity in the population are major strengths.

This study had some limitations. First, given that this study uses data from cross-sectional studies, results focused on associations and not causality [37]. The current lack of data to identify genes contributing to obesity in this study made it impossible for us to examine the biological pathways and how its interaction with environmental factors influenced the risk of obesity. Future studies may need to identify and examine the effect of obesity predisposing genes to determine such biological pathways. Additionally, the study focuses only on those who were 50 years and above and does not cover the entire population. Therefore, conclusions from this study is limited to the population of 50 years and above. Finally, although the data is representative of the population aged 50 years and over, the analysis omits observations with missing data (13% in Wave 1 and 4.8% in Wave 2) on variables such as weight, height and waist circumference. Hence, there is a chance for selection bias to be introduced that might have affected external validity. However, the analytical sample was large, and the use of the post-stratified persons’ weight supported the analysis.

## Conclusion

We found that although underweight has reduced in Ghanaian older adults, the prevalence of overweight among males and females, and obesity among females has increased. The Ghana NCDs management strategy 2012-2016 focused on reducing by 2% the overweight and obesity prevalence in females age 15-49years, neglecting those aged 50+ years. However, the exponential increase in the current estimates of obesity prevalence suggest the need for policy initiatives aimed at reducing overweight and obesity, especially among females aged 50+ years and the importance of advancing surveillance. This study also identifies key factors that could be targets of tailored prevention and intervention programs in improving the health and well-being of older adult populations in sub-Saharan Africa.

## DECLARATIONS

### Acknowledgement

We appreciate access to a preliminary version of SAGE Ghana Wave 2 data used for the analyses in this manuscript.

### Ethics and permissions

Data from SAGE Ghana Wave 2 was used for this study. WHO SAGE was approved by the WHO Ethics Review Committee (reference number RPC149) with local approval from the University of Ghana Medical School Ethics and Protocol Review Committee (Ghana). The necessary permission was obtained from the World Health Organization to use these data.

### Data Availability Statement

Data from SAGE Ghana Wave 2 was used for this study. WHO SAGE was approved by the WHO Ethics Review Committee (reference number RPC149) with local approval from the University of Ghana Medical School Ethics and Protocol Review Committee (Ghana). The necessary permission was obtained from the World Health Organization to use these data. All files were obtained from the World Health Organization Study on global AGEing and adult health (WHO-SAGE). Details on data can be found at http://www.who.int/healthinfo/sage/cohorts/en/. The authors used the GhanaINDDataW2 and GhanaHHDataW2. The codes for the measured weight and height used to calculate body mass index (BMI), as used in the data are q2506 for weight and q2507 for height. The authors confirm that they had no special access privileges to the data. Interested researchers will have to submit a licensed data request to WHO. Upon approval, the researchers will be granted access licensed dataset.

### Compliance with ethical standards

#### Funding

The authors received no specific funding for this work.

Prof Andrew J Palmer is funded by the Centre of Excellence in Population Ageing Research, Australian Research Council (CE170100005).

The National Heart Foundation of Australia Future Leader Fellowship (100849) supports Dr. Costan G Magnussen.

Dr. Lei Si is supported by a NHMRC Early Career Fellowship (Grant number: GNT1139826).

Dr Barbara de Graaff is funded by the Menzies Community Fellowship.

The Study on global AGEing and adult health (SAGE) Wave 2 was supported by WHO and the US National Institute on Aging’s Division of Behavioural and Social Science Research (BSR) through Interagency Agreements (OGHA 04034785; YA1323-08-CN-0020; Y1-AG-1005-01) with WHO. Financial and in-kind support has come from the University of Ghana’s Department of Community Health.

### Authors’ Contribution

STL, LS, CGM, BdG, and AJP formulated the initial research idea and designed the study. STL, LS, CGM, BdG, LB, and AJP analysed the data. STL, LS, CGM, BdG, GOB, LB and AJP wrote, reviewed and edited the manuscript. RBB, NM and PK reviewed and edited the manuscript.

